# Network-centered homeostasis through inhibition maintains hippocampal spatial map and cortical circuit function

**DOI:** 10.1101/2020.08.04.236042

**Authors:** Klara Kaleb, Victor Pedrosa, Claudia Clopath

## Abstract

Despite ongoing experiential change, neural activity maintains remarkable stability. Such stability is thought to be mediated by homeostatic plasticity and is deemed to be critical for normal neural function. However, what aspect of neural activity does homeostatic plasticity conserve, and how it still maintains the flexibility necessary for learning and memory, is not fully understood. Homeostatic plasticity is often studied in the context of neuron-centered control, where the deviations from the target activity for each individual neuron are suppressed. However, experimental studies suggest that there are additional, network-centered mechanisms. These may act through the inhibitory neurons, due to their dense network connectivity. Here we use a computational framework to study a potential mechanism for such homeostasis, using experimentally inspired, input-dependent inhibitory plasticity (IDIP). In a hippocampal CA1 spiking model, we show that IDIP in combination with place tuned input can explain the formation of active and silent place cells, as well as place cells remapping following optogenetic silencing of active place cells. Furthermore, we show that IDIP can also stabilise recurrent network dynamics, as well as preserve network firing rate heterogeneity and stimulus representation. Interestingly, in an associative memory task, IDIP facilitates persistent activity after memory encoding, in line with some experimental data. Hence, the establishment of global network balance with IDIP has diverse functional implications and may be able to explain experimental phenomena across different brain areas.

## Introduction

Although neural activity varies, presumably due to different demands of various cognitive functions, it is usually constrained to a small operational range of only a few Hertz. This regime may facilitate energy saving (Perrinet 2010), as well as optimal information processing (Laughlin 1981, Stemmler & Koch 1999). Furthermore, deviations from this range are often associated with pathological states, such as epilepsy, schizophrenia, and Fragile X syndrome (Dickman & Wondolowski 2013). Given that neural activity undergoes constant experiential change, there exists a requirement for active processes to maintain stability. Many such processes have been identified, such as synaptic scaling (Turrigiano et al. 1998, Desai et al. 2002, Turrigiano & Nelson 2004, Goel & Lee 2007, Glazewski et al. 2017), intrinsic plasticity (Desai et al. 1999, Gainey et al. 2018, Lambo & Turrigiano 2013, Maffei & Turrigiano 2008), meta-plasticity (Bienenstock et al. 1982, Kirkwood et al. 1996, Zenke et al. 2013, Frank et al. 2006), diffusive neuromodulation (Sweeney et al. 2015, Steinert et al. 2008, 2011, Naumann & Sprekeler 2020), structural plasticity (Yin & Yuan 2015, Gallinaro & Rotter 2018) and inhibitory plasticity (Woodin et al. 2003, Maffei et al. 2004, 2006, Chen et al. 2011, Vogels et al. 2011, Keck et al. 2011, van Versendaal et al. 2012, D’amour & Froemke 2015, Udakis et al. 2019, Clopath et al. 2016, Hennequin et al. 2017, Das et al. 2011, Haas et al. 2006). These altogether make up the term homeostatic plasticity and, although vastly different, they all act as a negative feedback mechanism that adjust the neural parameters to compensate for deviations from some set-point. Homeostatic plasticity is often studied in the context of neuron-centered control, as neurons return to their preferred level of activity after experimental manipulations that decrease (Hengen et al. 2013) or increase activity (Pacheco et al. 2019). There is also experimental evidence of network-centered homeostasis (Hirase et al. 2001, Slomowitz et al. 2015, Trouche et al. 2016), where instead of the individual neurons, the mean activity of the whole network is homeostatically maintained. However, computational studies of such mechanisms are few (Sweeney et al. 2015, Naumann & Sprekeler 2020) and thus they remain less well understood.

To illustrate network-centered homeostasis, we turn to the hippocampus. Spatial environments are known to be represented by a cognitive map, consisting of a subset of hippocampal pyramidal cells. These are called place cells, as they fire action potentials when the animal is in a specific location within the environment, their place fields (O’Keefe & Dostrovsky 1971, O’Keefe 1976, O’keefe & Nadel 1978, Wilson & McNaughton 1993). Optogenetic silencing of the CA1 pyramidal neurons encoding a place map of a familiar environment leads to rapid activation of previously silent CA1 pyramidal neurons (Trouche et al. 2016). This abrupt increase is followed by a slower, seconds-long activity change towards a stable level on par with that of the original place map (Trouche et al. 2016). Thus, an alternative place map transiently emerges while the original place cells are being silenced, and the spatial representation is homeostatically maintained. Furthermore, with repeated silencing, the alternative place map is consolidated over the original place map (Trouche et al. 2016). As neurons presumably have access only to their own activity, it is unclear how such a pertubation can be detected and compensated for at the network level.

One of the likely candidates to implement such network-centered homeostasis are the inhibitory neurons. CA1 inhibitory neurons are strongly connected to a large number of heterogeneously tuned CA1 pyramidal cells (Ali et al. 1998, Freund & Buzsaki 1996, Gulyás et al. 1999, Bezaire & Soltesz 2013, Csicsvari, Czurko, Hirase & Buzsáki 1998, English et al. 2017, Csicsvari, Hirase, Czurko & Buzsaki 1998), and exhibit broad spatial tuning (Grienberger et al. 2017). Thus, they can facilitate lateral inhibition between the pyramidal cells (Geisler et al. 2007) and as such, both sense and influence the activity of their local network. Hence, plasticity in the inhibitory circuitry could offer a highly efficient network activity homeostasis, which has previously been shown with neuron-centered plasticity (Vogels et al. 2011). However, there are indications that inhibitory plasticity can also act more globally. For instance, inhibitory postsynaptic potentials (iPSPs) have been shown to decrease when global, but not single neuron, spiking activity is suppressed (Hartman et al. 2006, Peng et al. 2010). Moreover, in the highly recurrent pyramidal cell networks of the neocortex, where inhibitory neurons are similarly interconnected with the rest of the network (Sohya et al. 2007, Niell & Stryker 2008, Kerlin et al. 2010, Ma et al. 2010, Zariwala et al. 2011, Znamenskiy et al. 2018, Wilson et al. 2017, Fino & Yuste 2011, Packer & Yuste 2011, Hofer et al. 2011, Bock et al. 2011, Pfeffer et al. 2013), inhibitory scaling was shown to be Ca2+/calmodulin-dependent protein kinase IV (CAMKIV) independent (Joseph & Turrigiano 2017), and thus may be decoupled from unique postsynaptic neuron activity. Furthermore, sensory deprivation studies across the primary cortices (Kuhlman et al. 2013, Barnes et al. 2015, Li et al. 2014, Gainey et al. 2018) have led to suggestions that inhibition could be broadly adjusted as a function of the activity of the circuit (Gainey & Feldman 2017). Inspired by all of these experimental findings, we hypothesize that the synaptic input to the inhibitory neurons could act as a proxy for the local network activity, and as such be used to adjust the level of inhibition appropriately.

In this work, we computationally study the properties and potential functionality of such novel inhibitory plasticity, which we term input-dependent inhibitory plasticity (IDIP). First, we show that such plasticity provides a mechanistic explanation for the emergence of active and silent place cells in a hippocampal CA1 network model. Our model also reproduces the previously unexplained experimental data of fast and reversible remapping following acute optogenetic place map silencing (Trouche et al. 2016), as well as the consolidation of the alternative place map following repeated silencing (Trouche et al. 2016). Furthermore, we show that IDIP in a cortical recurrent network model provides a form of rapid firing rate homeostasis while maintaining important network features, such as firing rate heterogeneity and persistent activity. Thus, we show that IDIP allows for accurate maintenance of neural representation while preserving flexibility important for neural coding.

## Results

### Input-dependent inhibitory plasticity (IDIP) rule as a homeostatic mechanism

Experimental studies suggest an existence of network-centered homeostasis (Hirase et al. 2001, Slomowitz et al. 2015, Trouche et al. 2016), where the mean firing rate of the network, rather than that of the individual neurons, is homeostatically maintained. For example, rapid homeostasis of place representation in the hippocampal CA1 region is observed after optogenetic silencing of a familiar place map, through the emergence of an alternative place map (Trouche et al. 2016). The likely candidates underlying such a mechanism are the inhibitory neurons, due to their dense interconnectivity with the surrounding pyramidal cells, both in the hippocampus (Ali et al. 1998, Freund & Buzsaki 1996, Gulyás et al. 1999, Bezaire & Soltesz 2013, Csicsvari, Czurko, Hirase & Buzsáki 1998, English et al. 2017, Csicsvari, Hirase, Czurko & Buzsaki 1998) and in the neocortex (Sohya et al. 2007, Niell & Stryker 2008, Kerlin et al. 2010, Ma et al. 2010, Zariwala et al. 2011, Znamenskiy et al. 2018, Wilson et al. 2017, Fino & Yuste 2011, Packer & Yuste 2011, Hofer et al. 2011, Bock et al. 2011, Pfeffer et al. 2013). Furthermore, rapid disinhibition has been shown to be the first step in circuit reorganization after experimental deprivation across the sensory cortices (Kuhlman et al. 2013, Barnes et al. 2015, Li et al. 2014, Gainey et al. 2018), leading to suggestions that inhibition could be broadly adjusted based on the activity of the circuit (Gainey & Feldman 2017). Such global regulation of network activity could be achieved through the modulation of inhibition as a function of the synaptic input the inhibitory neurons receive. To this end, we hypothesize a plasticity mechanism in which strong inputs onto the inhibitory neurons lead to strengthening of the inhibitory output, whereas weak inputs onto the inhibitory neurons lead to weakening of the inhibitory output. For simplicity, we choose to implement this input-dependent inhibitory plasticity (IDIP) rule by scaling the inhibitory synaptic weights as a function of the difference between the actual and the target synaptic input that each inhibitory neuron receives (Fig 1.A). Note that such cell-autonomous plasticity could also be implemented through the plasticity of excitability of the inhibitory neurons, which is not explored in this work.

**Figure 1.**
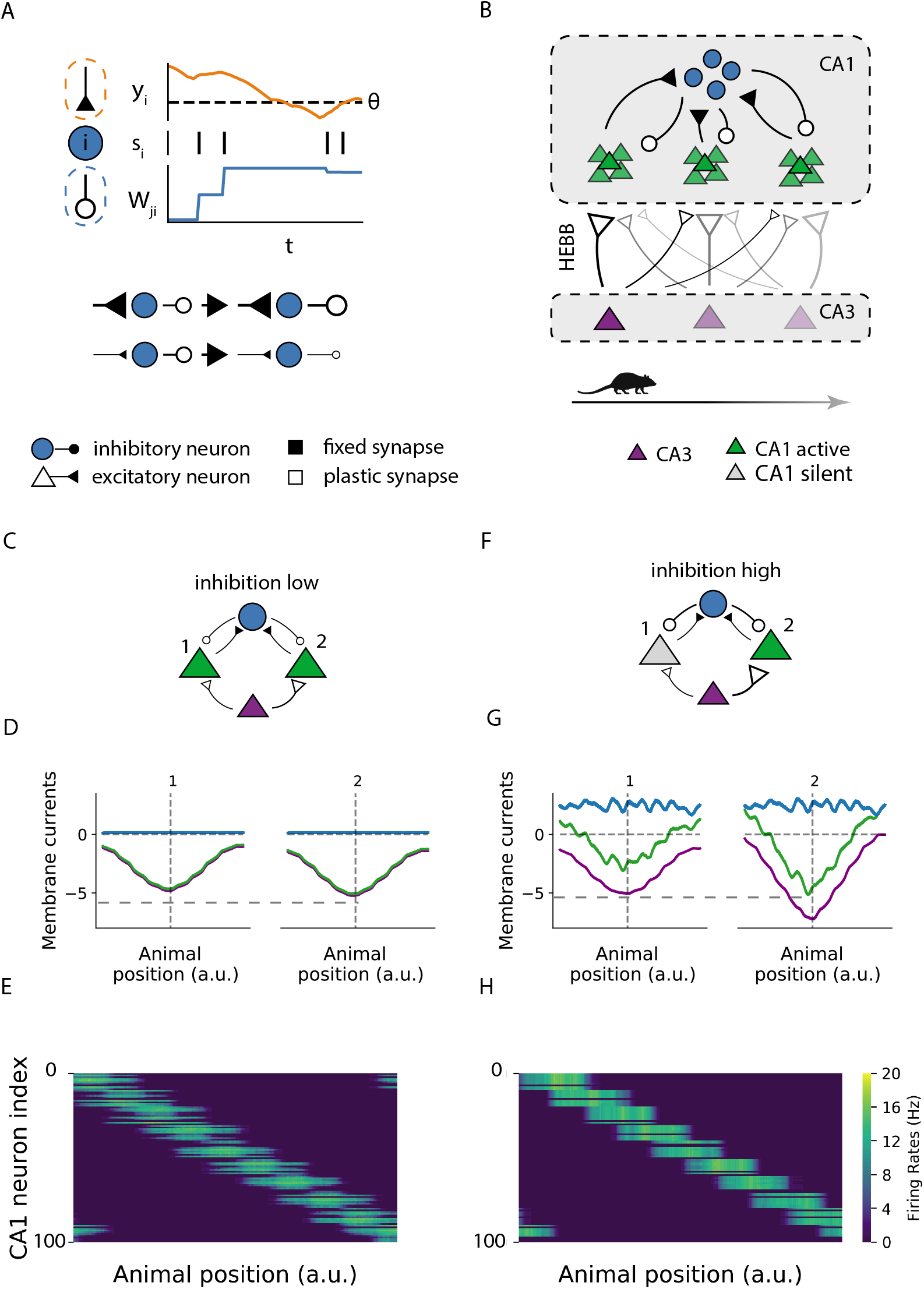
The IDIP rule enables emergence of active and silent place cells in a hippocampal network. **A.** Diagram of the inhibitory plasticity rule. An inhibitory neuron (blue) adjusts its projecting inhibitory synaptic weights based on the synaptic current it is receiving. Strong inputs onto inhibitory neurons lead to strengthening of inhibitory output weights, whereas weak inputs onto inhibitory neurons lead to weakening of inhibitory output weights. **B.** Hippocampal network diagram. The CA3 region consists of 10 excitatory neurons (purple triangles). Each receives unique place-dependent external current, representing the location of a simulated mouse on a 1D annular track with equally spaced place fields. CA3 neurons project to 100 CA1 excitatory neurons (green triangles). These are split in 10 equally sized groups, with neurons in each group tuned to the same place field. CA1 excitatory neurons project to CA1 inhibitory neurons (blue circles), which in turn project back to the CA1 excitatory neurons. The excitatory synapses from the CA3 to the CA1 are plastic under Hebbian plasticity and the inhibitory synapses in the CA1 are plastic under IDIP. **C-E** State of the network during the first lap, before any learning. **C.** A diagram of a sample microcircuit found in the network B, consisting of 2 CA1 excitatory neurons (1 and 2) with the same spatial tuning, but with neuron 2 having slightly larger CA3 to CA1 synaptic strengths. **D.** The excitatory (purple), inhibitory (blue) and net (green) currents received by neurons 1 and 2 during the first lap on the track. The higher peak net current received by neuron 2 reflects its higher place field tuning compared to neuron 1. **E.** The firing rates of all the excitatory neurons in the CA1 network (y-axis) during the first lap on the simulated track (x-axis). All the place cells are active but with varying amplitudes, reflecting differences in their place tuning strengths. **F-H** Same as in B-D, but after inhibitory and excitatory learning (100 laps). **F.** The same microcircuit diagram as in C but after learning. The CA3 to CA1 excitatory synaptic plasticity leads to increased activity of both neurons. Due to the differences in their tuning, the excitatory synapses to neuron 2 are potentiatied more. At the same time, increased activity in the CA1 excitatory neurons increases input to the inhibitory neurons, and thus the inhibitory synaptic weights are increased as well. This increased inhibition eventually completely overcomes excitation in the less active neuron 1 and hence it becomes a silent place cell (grey). **G.** Excitatory (orange), inhibitory (blue) and net (green) currents received by neurons 1 and 2 during the last lap on the track. The difference in the peak net currents from the first lap **D** is amplified with learning. **H.** The firing rates of all the excitatory neurons in the CA1 network (y-axis) during the last lap on the simulated track (x-axis). The place cells have a well defined place field and a subset of cells within each place tuned group have become silent.

### The IDIP rule allows for the emergence of active and silent place cells in a hippocampal network

To assess whether IDIP can lead to the emergence of active and silent place cells, we build a hippocampal network model of leaky integrate-and-fire neurons. The network model consists of the CA3 and CA1 region (Fig 1.B). Each pyramidal cell in the CA3 region receives unique place tuned external current (Fig S1.A-B), representing the location of a simulated mouse on a 1D annular track with equally spaced place fields. The CA3 neurons project to the CA1 excitatory neurons, which are divided into equally sized groups, with neurons in each group tuned to the same place field. As the CA1 excitatory neurons are poorly recurrently connected (Deuchars & Thomson 1996, Thomson & Radpour 1991), in our simulations we assume that there are no recurrent connections between them. The CA1 excitatory neurons project to the CA1 inhibitory neurons and the CA1 inhibitory neurons project back to the CA1 excitatory neurons. The CA1 inhibitory neurons have no spatial tuning (Fig S2.A), in agreement with experiments (Dupret et al. 2013, Grienberger et al. 2017). The excitatory synapses between the CA3 and CA1 excitatory neurons and the inhibitory synapses to the CA1 excitatory neurons are made plastic with Hebbian plasticity and the IDIP rule respectively.

We first wanted to test whether IDIP could allow for the emergence of both active and silent place cells in the CA1 region. The experiments suggest that the cells which later become active or silent are differentiable even before the first exploration of the environment (Epsztein et al. 2011). We incorporate this in our model by introducing variability in the amplitude of the place tuning of the CA1 excitatory neurons belonging to the same group (Fig 1.C-D, Fig S3.A-B). As the initial inhibitory synaptic weights are set to low values, all the CA1 excitatory neurons are active, covering the whole environment (Fig 1.E). We then let our simulated mouse complete 100 laps on the track, which we term the exploration phase. During this phase, the CA1 excitatory neurons that are highly place tuned are potentiated more. Through activity, their tuning also increases in a positive feedback loop characteristic to Hebbian learning (Fig 1.F-G, Fig S3.C-D). This increases the synaptic input to the CA1 inhibitory neurons, and hence increases the inhibitory synaptic weights through IDIP (Fig S2.B). As the CA1 excitatory neurons with lower place tuning are less active, they are unable to escape the increasing lateral inhibition. Thus stable active and silent place cells form within each CA1 group (Fig 1.H, Fig S3.E-F). The absence of the target firing rate for individual excitatory neurons is essential for this. Hence, we show that the IDIP rule together with Hebbian plasticity and place tuned inputs can explain the formation of active and silent place cells in a hippocampal network model.

### The IDIP rule enables rapid homeostatic remapping during active place cells silencing

To assess whether the proposed IDIP rule can facilitate rapid place cell remapping, we perform a silencing protocol as in Trouche et al. (2016) (Fig 2.A). After the exploration phase, we tag all the established active place cells and silence them. In the networks with (Fig 2.B) and without (Fig. 2.F) IDIP, such silencing leads to a decrease in the excitatory drive to the CA1 inhibitory neurons and subsequently a rapid decrease in the inhibition to the previously silent place cells, which then form an alternative place map. However, the degree of the alternative place map activation will depend on inhibitory plasticity. In the networks with IDIP during the silencing (Fig 2.B), the dynamics of the alternative place map activation are similar to that observed experimentally (Fig 2.C) (Trouche et al. 2016). The firing rates of the alternative place cells are first increased almost immediately after silencing onset (fast phase) and are further increased within the next 1-2 seconds (slow phase) (Fig 2.D). Our model allows us to decompose the two phases. The fast phase is due to the rapid decrease in CA1 inhibitory neuron firing rate (Fig S4.A), as the originally active place cells, which were driving their activity, are no longer active. Thus the alternative place map emerges, but at a lower firing rate than the original place map, due to the place cells forming the alternative place map being less sharply place-tuned than the place cells that formed the original place map. In the networks without IDIP during the silencing (Fig 2.F), the activity of the alternative place map does not progress beyond this phase (Fig 2.G-H). However, in the networks with IDIP during the silencing, a second, slower phase of alternative map activation occurs as the level of inhibition gets adjusted to the lower activity and thus lower synaptic input from the alternative place cells (Fig S4.B). Therefore the final activity of the alternative place map matches that of the original place map (Fig 4.E). Hence, the IDIP rule acts as a homeostatic mechanism to maintain network activity during acute silencing.

**Figure 2.**
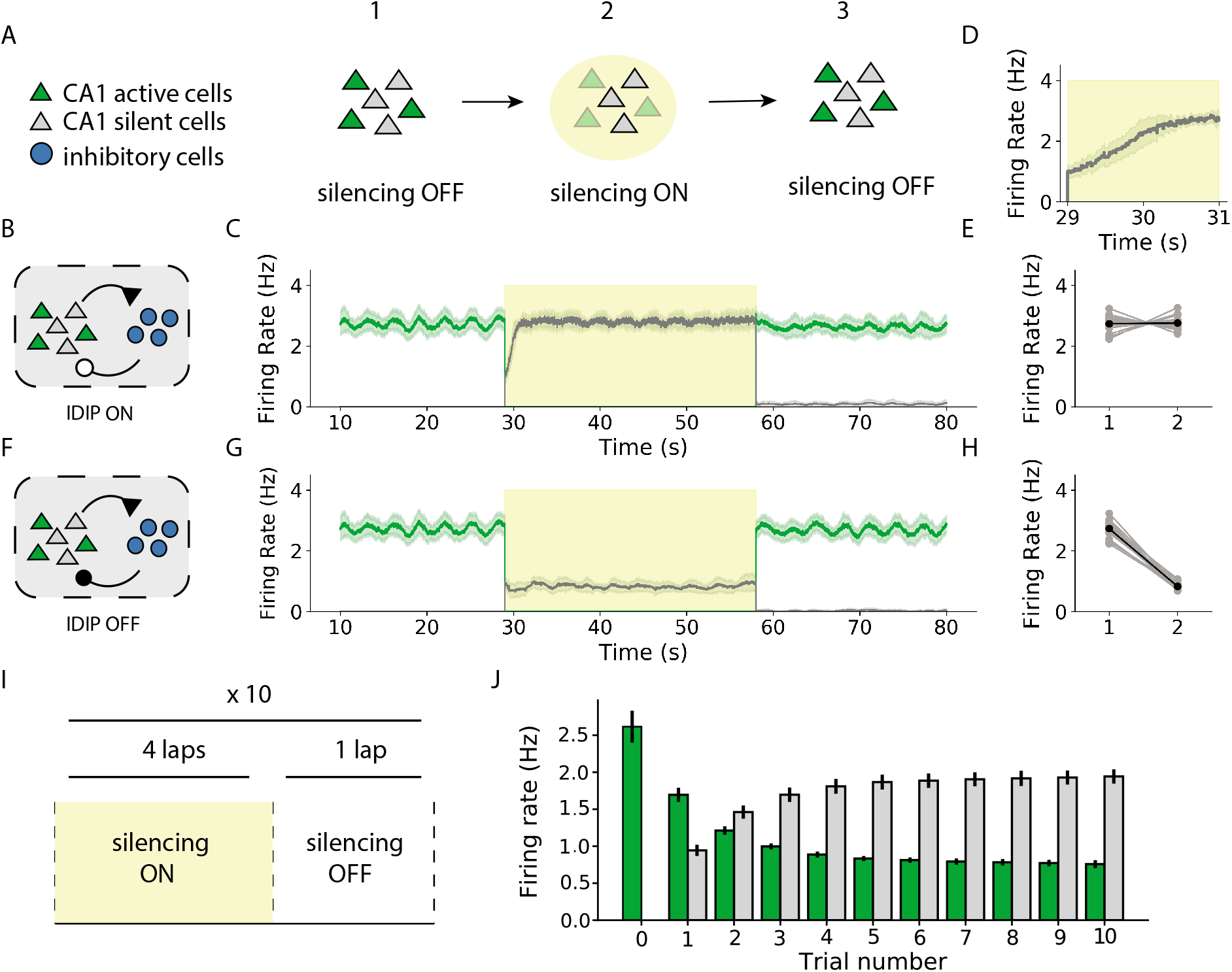
Place cells silencing leads to a rapid emergence of an alternative place map. **A.** Silencing protocol similar to Trouche et al. (2016). All the active cells (green) are tagged (1), silenced for one lap (2) and released in the subsequent lap (3). **B-E** Silencing in the network following IDIP. **B.** Network diagram after the exploration phase (100 laps). **C.** Mean firing rate of the network just before (1), during (2), and after silencing (3). During silencing (2) there is rapid activation of an alternative place map, consisting of the previously silent cells (grey). After silencing (3), the original map reactivates (green). **D.** Alternative place map activation just after silencing. **E.** The mean firing rate of the place map before (1) and during (2) silencing. Gray circles indicate individual trials. Black circles indicate average over all 20 trials. **F-H** Same as in **B-E** but with inhibitory plasticity turned off during silencing (2). **F.** Network diagram after inhbitory and excitatory learning (lap 100) with IDIP turned off during silencing (2). **G.** Without IDIP during the silencing (2) there is low alternative place cell activation. **H.** Without IDIP during the silencing (2) the mean firing rate of the place map decreases. Gray circles indicate individual trials. Black circles indicate average over all 20 trials. **I** Consolidation protocol similar to Trouche et al. (2016). The established place map is silenced for 4 consecutive laps, and then released for a single testing lap. This is repeated for 10 trials. **J.** Mean firing rate of the active (green) and silent (grey) place map during the testing laps at each trial. Trial 0 corresponds to the activity of the network just after the exploration phase.

**Figure 3.**
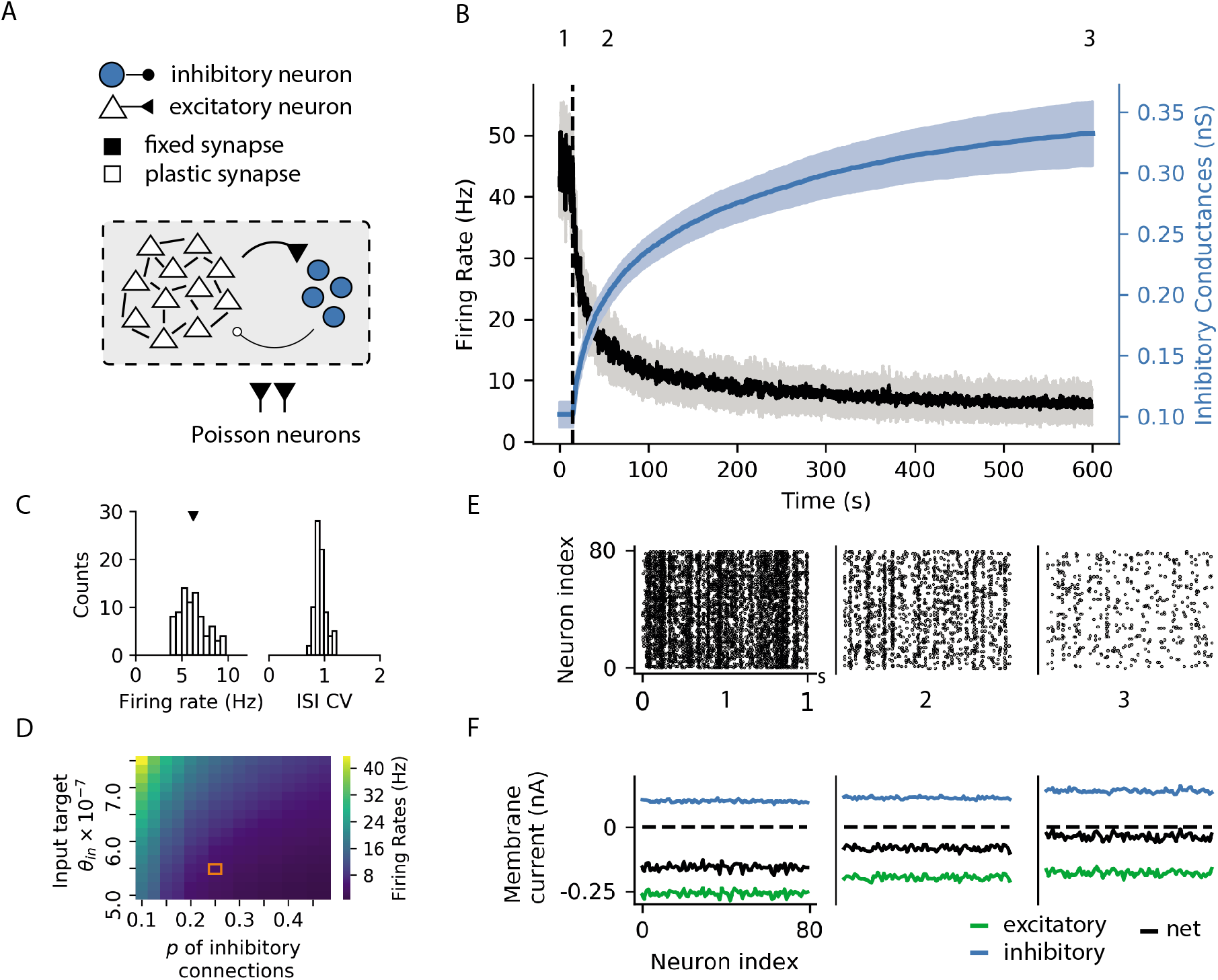
IDIP establishes global E/I balance in recurrent networks. **A.** A recurrent network of 80 excitatory (white triangles) and 20 inhibitory (blue circles) leaky integrate-and-fire neurons receiving external input from a pool of 100 Poisson excitatory neurons. Only the inhibitory synaptic weights are made plastic with IDIP. **B.** The evolution of the mean excitatory firing rate (black) and the mean inhibitory weight (blue) of the network. Shaded regions correspond to the standard deviation of individual units. IDIP is turned on at 15s (dashed vertical line). Activity of the network decreases as inhibitory weights increase. **C.** The excitatory firing rate stabilises to a mean value of 6.2 Hz (left, black triangle). The network displays an asynchronous firing pattern (right). **D.** The range of mean network firing rates IDIP can support, with varying input target value (y-axis) and inhibitory connectivity (x-axis). The orange square marks the parameters used in our simulations. **E.** Spike raster plots at the time points marked in **B**. The network progresses from high synchronous to low asynchronous firing following the IDIP rule. **F.** Mean membrane currents as a function of excitatory neuron index at the time points marked in **B**.

**Figure 4.**
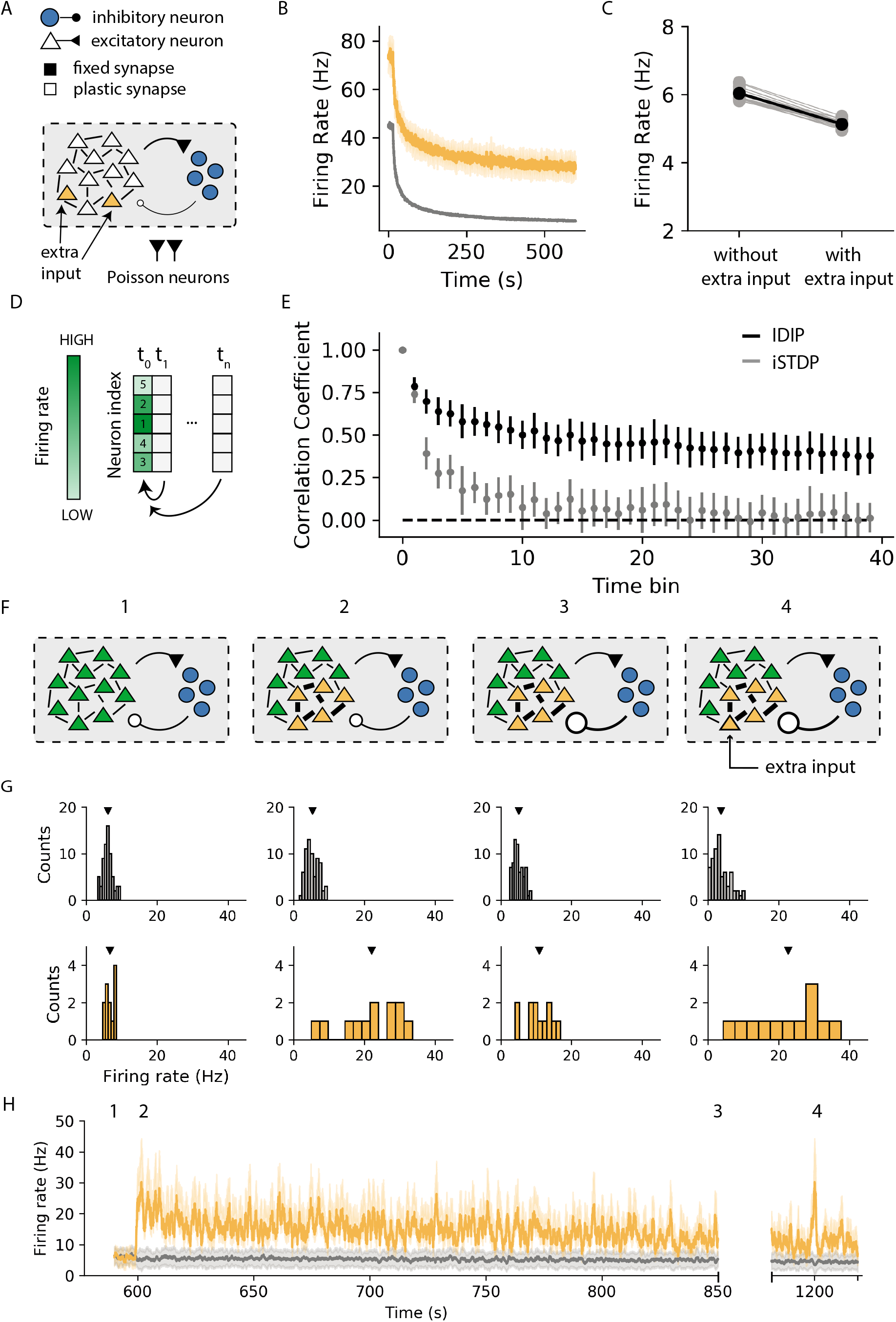
IDIP rule enables maintenance of neural representations and memory trace persistence. **A-C.** Perfomance of the IDIP rule in networks with heterogeneous activity. **A.** The same network as in Fig 3.A, but with a subset of neurons receiving increased external input. **B.** The average firing rates of the networks with increased input as in A. Bold lines denote the average and the shaded area indicated standard deviation over 20 trials. The increased activity relative to the rest of the network is sustained after inhibitory learning with IDIP. **C.** The average firing rate of the networks without (left) and with (right) input heterogeneity. Gray circles indicate individual trials and the black circles indicate average over all 20 trials. **D-E.** Performance of IDIP and iSTDP (Vogels et al. 2011) in a network representation task. **D.** Network representation task diagram. We calculate the preservation of the correlation with the initial rank of firing rates during the entire course of the simulation. **E.** The Spearman rank correlation coefficient for the network following IDIP (black) and iSTDP (grey). Error bars correspond to standard deviation over 20 trials. The horizontal dashed line indicates no correlation. **F-H.** Perfomance of the IDIP rule in an associative memory task. **F.** Associative memory task protocol. Following network stabilization (1), we increase recurrent excitatory connections between a subset of neurons (2). This leads to increase in inhibition (3). The encoded assembly is recalled by increasing external input to a subset of neurons within the pattern (4). **G.** Histograms of average firing rates of the assembly (orange) and the rest of the network (grey) at the marked task time points **F.** The black triangles indicate the mean firing rates. **H.** The evolution of the firing rates of the assembly (orange) and the rest of the network (grey) during the memory task. The shaded area indicates standard deviation over all units in each network subset.

When we turn silencing off, the original place map re-emerges (Fig 2.C), in agreement with experiments (Trouche et al. 2016). However, repeating the silencing protocol (Fig 2.I) consolidates the alternative place map (Fig 2.J), also in agreement with experiments (Trouche et al. 2016). In our model this happens through the gradual activity-dependent Hebbian plasticity in the CA3 to CA1 excitatory synapses (Fig S5.A). Therefore, our model suggests that synaptic plasticity of feedforward inputs onto the CA1 pyramidal cells is a good candidate for the mechanism underlying place map consolidation.

### The IDIP rule establishes global E/I balance in recurrent networks

The inhibitory neurons also feature dense connectivity in the neocortical circuits (Sohya et al. 2007, Niell & Stryker 2008, Kerlin et al. 2010, Ma et al. 2010, Zariwala et al. 2011, Znamenskiy et al. 2018, Wilson et al. 2017, Fino & Yuste 2011, Packer & Yuste 2011, Hofer et al. 2011, Bock et al. 2011, Pfeffer et al. 2013) and there is some evidence for a network-centered control of cortical activity through inhibition (Joseph & Turrigiano 2017, Gainey & Feldman 2017, Gainey et al. 2018). To assess whether the proposed IDIP learning rule can homeostatically regulate such circuits, we simulate a cortical network of leaky integrate-and-fire neurons with sparse recurrent connectivity (Fig 3.A). Each neuron in the network receives large external excitatory input and the initial inhibitory synaptic weights are set to low values. Thus, without any plasticity, the network exhibits pathologically high activity (Fig 3.B). Due to the high activity of the network, the inhibitory neurons initially receive very large excitatory synaptic input, and thus the IDIP rule increases the inhibitory synaptic weights (Fig 3.B - IDIP turned on at 15s, Fig S6.A). The network, therefore, progresses from a high synchronous to a low asynchronous firing following the IDIP rule (Fig 3.E). The excitatory firing rates reach a more physiological regime (Fig 3.B, Fig S6.B), with a reasonable firing rate distribution and irregular firing rate dynamics (Fig 3.C). Inhibitory neurons in the network also exhibit a diversity of firing rates and irregular firing rate dynamics (Fig S7.A-E). Hence, IDIP can homeostatically regulate network-wide activity.

The IDIP rule proposed does not impose a unique target firing rate for each neuron in the network. Instead, it controls the mean firing rate across the whole network. The final variability in the net current received by each individual excitatory neuron after inhibitory learning (Fig 3.F) results in a diversity of neuronal firing rates (Fig 3.C). This is in contrast with the network following a neuron-centered rule, such as inhibitory spike-timing dependent plasticity (iSTDP) (Vogels et al. 2011), which effectively sets a target firing rate for the excitatory neurons (Fig S8.A-E). To assess the range of the firing rates the IDIP rule can support in a recurrent network, we simulate the network for various values of the inhibitory target input. Higher values of the inhibitory target input lead to higher network activity (Fig 3.D, y-axis). Hence, the activity of the network following the IDIP rule depends on the target input of the inhibitory neurons. Altogether, this means that IDIP establishes a global, rather than detailed, network excitatory (E)/ inhibitory (I) balance while allowing for network diversity.

### The IDIP rule enables maintenance of neural representations

We wanted to assess whether neural representation could be maintained in the networks following the IDIP rule. To assess whether individual neurons can remain highly active, we increase the external drive to a subset of neurons in the network (Fig 4.A). The selected subset sustains a higher firing rate relative to the rest of the network following inhibitory learning with IDIP (Fig 4.B, S9.A). This effect is mediated by the highly active neural subset monopolizing the input to the inhibitory neurons, leading to greater inhibition to the rest of the network. Thus the rest of the network compensates for this increased activity, as seen in a slight decrease in the mean network firing rate (Fig 4.C). As expected, the deviations from the target excitatory neuron firing rate are suppressed in the network following iSTDP (Vogels et al. 2011) (Fig S9.B). Hence, IDIP can control the recurrent network activity while preserving activity heterogeneity.

To assess whether the relative firing rates between neurons can be conserved across the whole neural population, we rank the firing rates of the neurons at the beginning of our simulations, before any inhibitory plasticity (Fig 4.D). We then measure whether this rank is maintained over time following inhibitory plasticity (Fig 4.D). In networks following IDIP, the firing rate rank correlation is largely conserved after inhibitory learning (Fig 4.E, Fig S9.C). We compare this with the network following iSTDP (Vogels et al. 2011), where the firing rate rank correlation is mostly lost, as all neurons fire at similar firing rates at the end of the simulations (Fig 4.E, Fig S9.D). Hence, IDIP can preserve heterogeneity in firing rates, both at a single neuron level and across the whole network.

### The IDIP rule enables memory trace persistence and recall

We also assess the performance of the network following IDIP in a simple associative memory task. Once the network activity stabilises, we encode a memory by increasing the recurrent excitatory connections within a pattern in the network (Fig 4.F), as previously done in networks following iSTDP (Vogels et al. (2011), reproduced in Fig S10.C-D). After encoding, the neurons recruited to the memory pattern exhibit sustained activity which is higher than the rest of the network, even after IDIP has converged (Fig 4.G-H, Fig S10.A). This is in contrast to the networks following iSTDP (Vogels et al. 2011), in which the activity of the memory ensemble becomes indistinguishable from the rest of the network at convergence (Fig S10.C-D). Such persistent activity of the memory cells has been reported in some experiments (Yassin et al. (2010), Ghandour et al. (2019), but see Barron et al. (2017)).

We then test whether we can recall the pattern, given that IDIP made inhibition stronger in the network. By increasing the external current to some units in the pattern, the memory can be recalled (Fig 4.G-H, Fig S10.B). Performing the same protocol in a network following iSTDP (Vogels et al. 2011) shows that a lower number of memory cells are re-activated with recall (Fig S10.C). Thus recall in the network following IDIP has higher fidelity. This is in line with experiments (Pignatelli et al. 2019), where increased activity of the memory cells facilitates greater pattern completion. Hence, the IDIP rule allows for sustained activity of memory cells after encoding, as well as faithful memory recall from a partial cue.

In summary, we show that our proposed inhibitory plasticity rule, IDIP, can homeostatically regulate activity in models of hippocampal and cortical networks. Importantly, using IDIP we are able to reproduce previously unexplained experimental findings in the hippocampus (Trouche et al. 2016) and propose its potential functional implications in the recurrent cortical networks.

## Discussion

Although the study of neural homeostasis is frequently neuron-centered, there is some evidence that network-centered mechanisms are also at play (Hirase et al. 2001, Slomowitz et al. 2015, Trouche et al. 2016). One of the candidates proposed for the implementation of such homeostasis are the inhibitory neurons, due to their dense connectivity, both in the hippocampus (Ali et al. 1998, Freund & Buzsaki 1996, Gulyás et al. 1999, Bezaire & Soltesz 2013, Csicsvari, Czurko, Hirase & Buzsáki 1998, English et al. 2017, Csicsvari, Hirase, Czurko & Buzsaki 1998), as well as in the neocortex (Sohya et al. 2007, Niell & Stryker 2008, Kerlin et al. 2010, Ma et al. 2010, Zariwala et al. 2011, Znamenskiy et al. 2018, Wilson et al. 2017, Fino & Yuste 2011, Packer & Yuste 2011, Hofer et al. 2011, Bock et al. 2011, Pfeffer et al. 2013). In this work, we take inspiration from the experimental data and hypothesize that network homeostasis could be achieved through input-dependent inhibitory plasticity (IDIP), in which the inhibition is adjusted as a function of the synaptic input the inhibitory neurons receive. We show that in a hippocampal CA1 network model, IDIP can provide a mechanistic circuit understanding and reproduce experimental data in which place cells are optogenetically silenced, unmasking a previously silent place map (Trouche et al. 2016). Furthermore, we show that such a learning rule can also homeostatically regulate the activity of a recurrent neural network, while preserving flexibility important for neural coding. Altogether, our results suggest that network activity homeostasis following external manipulation or endogenous changes could share a common underlying mechanism.

In contrast to the commonly used neuron-centered inhibitory plasticity (Vogels et al. 2011), the IDIP rule features an absence of a target firing rate for each excitatory neuron. Instead, with the IDIP rule, the level of inhibition is modulated to maintain the mean firing rate of the whole network. The importance of this is seen in our data-driven model of the CA1 network, where IDIP allows for emergence of both active and silent place cells. In our model, these emerge as the neurons with higher activity dominate the input to the inhibitory neurons, and thus the recruitment of lateral inhibition. Conversely, the less active cells become silent. Thus, there is an amplification of slight differences in activity through scaling of the tonic inhibition, indiscriminately to all cells. Such competition for the recruitment of inhibition has previously been suggested to shape hippocampal assemblies (Buzsáki 2010). Furthermore, silencing of an established place map induces rapid compensatory adjustment of inhibition, and thus an emergence of an alternative place map (Fig 2.C), as reported experimentally (Trouche et al. 2016). We show that non-plastic inhibition is not consistent with the experimental data (Fig 2.G). Finally, we show that repeated silencing leads to alternative place map consolidation (Fig 2.J), as reported experimentally (Trouche et al. 2016). In our model, this occurs through gradual changes in the CA3 to CA1 excitatory synaptic weights (Fig S5), facilitated by disinhibition during each silencing lap (Fig S4.A). Thus, through time, the original place map gets destabilised, which has previously been suggested as a necessary condition for remapping (Schoenenberger et al. 2016). Hence, network-centered homeostasis through IDIP provides a possible explanation for the previously unexplained experimental phenomena in the hippocampus.

We also assess whether IDIP can homeostatically regulate dynamics in the recurrent networks typical of the neocortex. Due to the absence of a firing rate set-point in the networks following IDIP, there is diversity in the neuronal firing rates within the recurrent network after inhibitory learning (Fig 3.C). The resulting firing rate distribution is consistent with the broad and heavy-tailed distribution of firing rates observed experimentally (Wohrer et al. 2013), with some neurons more active than others (Mizuseki & Buzsáki 2013). Such range of activity is thought to be optimal for the information storage in the brain (Laughlin 1981) and enables linear network responses over a broad range of inputs (Sweeney et al. 2015). In our networks, highly active neurons dominate synaptic input to the inhibitory neurons, recruit inhibition more and thus inhibit other neurons more. Hence with the IDIP rule, highly active neurons can remain so as long as the rest of the network can compensate for it, providing a more flexible control. We illustrate this when we increase the external inputs to some neurons and they remain consistently more active than the rest of the network (Fig 4.B). Furthermore, we show the IDIP rule can also extend this stability of the firing rate rank across the whole neural population (Fig 4.E). This is not the case with inhibitory spike-timing dependent plasticity (iSTDP) (Vogels et al. 2011), as the ranks get shuffled following inhibitory learning, due to the unique firing rate set-point constraint (Fig 4.E). Hence, the IDIP model can maintain the population firing rate around a target value while preserving neural representation. Importantly, the neural activity diversity is not imposed but emerges due to random network structure. Although such population heterogeneity can be imposed, this has been previously shown to lead to limited network responsiveness (Sweeney et al. 2015). Hence, IDIP provides a global control of recurrent network dynamics, without a firing rate set-point for each neuron, which allows for flexibility and thus conservation of network representation.

Additionally, in an associative memory task, we show that the networks following the IDIP rule can sustain memory encoding, without disturbing the rest of the network (Fig 4.F-G). This has previously also been shown with iSTDP (Vogels et al. 2011). However, the key difference is that networks following the IDIP rule exhibit persistent activity of assemblies after encoding (Fig 4.G-H), unlike iSTDP (Fig S10.C-D), as well as higher fidelity of recall (Fig 4.G-H). Persistent activity of the memory cells has been reported in some experiments (Yassin et al. (2010), Ghandour et al. (2019), but see Barron et al. (2017)) and is thought to be involved in memory bindings via Hebbian plasticity, as the increased activity further strengthens the memory trace (Han et al. 2007) and facilitates greater recall (Tonegawa 2019). Thus, the global network control provided by IDIP may have important functional implications in memory processing and recall.

As highlighted throughout this work, dense connectivity of the inhibitory neurons to their local networks makes them an ideal candidate for network-centered homeostasis. However, more sparsely connected networks would lead the inhibitory neurons to sense a subset of the network and potentially affect a different subset. Thus the final network firing rate would be less constrained (Fig 3.D, Fig S11). This suggests that the network activity is differentially modulated by the IDIP rule depending on the network architecture. In the limit of very sparse networks, IDIP may not even act as a homeostatic mechanism and winner-take-all dynamics may dominate (Fig S12). This may be relevant in the prevention of redundancy or in contrast enhancement (Hartline & Ratliff 1957). Furthermore, inhibitory connectivity is known to vary within (Meyer et al. 2011) and between (Tamamaki & Tomioka 2010) brain areas. Thus, IDIP rule would have different functional consequences in different brain regions.

The basis of our proposed IDIP rule is that the inhibitory neurons, presumably fast-spiking parvalbumin (PV+) cells, can sense and affect the network activity through their input and output synaptic connections respectively. Input integration over the timescale used in our model could be mediated through N-methyl-D-aspartate receptors (NMDARs), which are known to enhance the neuronal computational repertoire (Gasparini & Magee 2006, Losonczy & Magee 2006, Poirazi & Mel 2001, Stuart & Spruston 2015, Poirazi & Papoutsi 2020). In the hippocampus, NMDAR knockouts in the PV+ cells disrupt spatial representation (Korotkova et al. 2010) and NMDAR were also shown to be preferentially localized at feedback synaptic inputs to the PV+ cells (Le Roux et al. 2013). NMDAR have also recently been shown to facilitate supralinear dendritic integration in the PV+ cells (Cornford et al. 2019), a model of which also shows winner-take-all dynamics between distinct assemblies (Cornford et al. 2019). Furthermore, in the prefrontal cortex, application of NMDAR antagonists causes disinhibition (Homayoun & Moghaddam 2007, Jackson et al. 2004, Zhang et al. 2008), initially caused by decrease in the inhibitory neuron activity (Homayoun & Moghaddam 2007), possibly via changes in their excitability, with prolonged exposures modulating the synthesis of gamma-Aminobutyric acid (GABA) (Zhang et al. 2008). Inhibitory output has also been shown to be modulated by activity, through modulation of rate limiting enzymes of GABA synthesis (Lau & Murthy 2012) or GABA transporters (De Gois et al. 2005). Such plasticity mechanisms have also recently been reported in the chandelier cells (Pan-Vazquez et al. 2020) and striatial inhibitory neurons (Paraskevopoulou et al. 2019) during development. Altogether, these studies indicate that our model of tonic inhibition modulated as a function of synaptic input to the inhibitory neurons is a biologically plausible mechanism.

Finally, other distinct solutions have been proposed for network-centered homeostasis, such as diffusive neuromodulation via nitric oxide (Sweeney et al. 2015) and GABA spillover (Naumann & Sprekeler 2020). It is very likely that they all coexist, alongside other, neuron-centered homeostatic mechanisms. Each homeostatic mechanism may control distinct aspects of neural function, as shown in a recent computational study (Wu et al. 2019). Hence, it would be of future interest to study the IDIP rule in conjunction with other homeostatic plasticity rules, such as synaptic scaling (Turrigiano et al. 1998, Desai et al. 2002, Turrigiano & Nelson 2004, Goel & Lee 2007, Glazewski et al. 2017), intrinsic plasticity (Desai et al. 1999, Gainey et al. 2018, Lambo & Turrigiano 2013, Maffei & Turrigiano 2008), metaplasticity (Bienenstock et al. 1982, Kirkwood et al. 1996, Zenke et al. 2013, Frank et al. 2006), diffusive neuromodulation (Sweeney et al. 2015, Steinert et al. 2008, 2011, Naumann & Sprekeler 2020), structural plasticity (Yin & Yuan 2015, Gallinaro & Rotter 2018) and inhibitory plasticity (Woodin et al. 2003, Maffei et al. 2004, 2006, Keck et al. 2011, Chen et al. 2011, Vogels et al. 2011, van Versendaal et al. 2012, D’amour & Froemke 2015, Udakis et al. 2019, Clopath et al. 2016, Hennequin et al. 2017, Das et al. 2011, Haas et al. 2006), as their interaction may increase the compensatory repertoire of the networks and endow them with non-trivial emergent properties.

## Methods

### Neuron Model

We use the single compartment leaky integrate-and-fire neuron model in our simulations. The model is defined by a resting membrane potential *V_REST_* and membrane time constant *τ_m_*. If the membrane potential surpasses the set threshold potential *θ_m_*, it fires a spike and its membrane potential is reset back to *V_REST_*. It then enters the refractory period *t_ref_*, during which it cannot be stimulated.

The sub-threshold membrane voltage *V_i_* of neuron *i* follows:

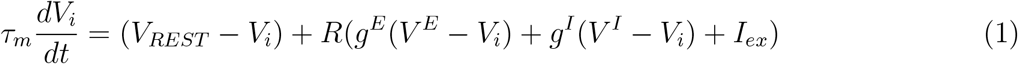

where *R* is the membrane resistance, *g_E/I_* are the excitatory and inhibitory synaptic conductances, *V^E/I^* are excitatory and inhibitory reversal potentials and *I_ex_* is any other externally supplied current. If neuron i receives input from neuron *j*, the corresponding synaptic conductance *g_ij_* is follows:

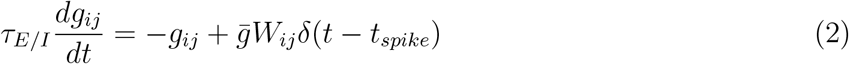

where *τ_E/I_* is the synaptic time constant, *W_ij_* is the synaptic weight, 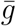 is a basic unit of synaptic conductance and *t_spike_* is the time of the presynaptic spike.

### Input-dependent inhibitory plasticity (IDIP)

The inhibitory plasticity depends on the input current *y_i_* received by each inhibitory neuron *i* over time.

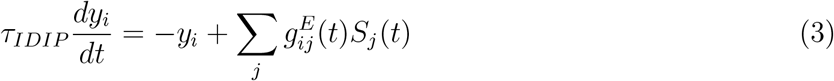

where *τ_IDIP_* is the input current time constant, 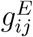 is the excitatory synaptic conductance of the synapse from an excitatory neuron *j* to an inhibitory neuron *i* and *S_j_*(*t*) is the spike train of the excitatory neuron *j*. Hence the second term is the sum of all of the excitatory input currents to the inhibitory neuron *i* at time *t*. When the inhibitory neuron i spikes, all the inhibitory synapses projecting from the presynaptic inhibitory neuron *i* to the postsynaptic excitatory neuron *j* are adjusted with respect to *y_i_* as:

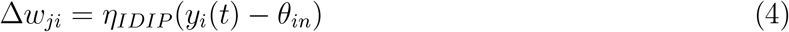

where *η_IDIP_* is the inhibitory learning rate and *θ_in_* is the constant target input for each inhibitory neuron. In our simulations, *θ_in_* is the same for all inhibitory neurons.

### Hippocampal Network Model

We simulate a CA1 network consisting of *N_E_* excitatory neurons and *N_I_* inhibitory neurons. There is full and bidirectional connectivity between the excitatory and inhibitory neurons. The initial synaptic weights from the inhibitory to the excitatory neurons (*W_IE_*) are 200 times weaker than the synaptic weights from the excitatory to the inhibitory neurons (*W_EI_*). The initial values of the inhibitory synaptic weights are set to ten times weaker than the excitatory weights. The CA1 excitatory neurons are divided in *N_g_* equally sized groups.

Each excitatory neuron in the CA1 network is connected to *N*_*CA*3_ CA3 neurons. The excitatory synaptic weights from the CA3 neurons to the CA1 excitatory neurons are made plastic with classical Hebbian plasticity and the inhibitory synaptic weights from the CA1 inhibitory neurons are made plastic with IDIP.

#### Position-modulated inputs

We model a mouse traversing a 1D annular track with *N_pc_* equally spaced place fields. We assume the mouse moves with a constant speed and takes 3 seconds to move from a centre of one place field to a centre of the subsequent place field.

Each CA3 neuron receives input from one unique place field in the form of an external current *I_ex_*. The tuning curve *T* with place field centered at *p*_0_ is defined as:

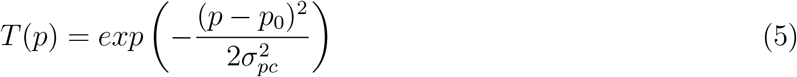

where *p* is the animal’s position and *σ_pre_* is the tuning width.

The place field current *I_ex_* supplied to a CA3 neuron is then:

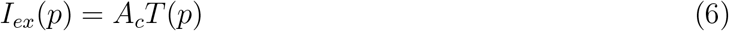

where *A_c_* is the current amplitude.

Each CA1 group is tuned to a single unique CA3 neuron, with the same tuning curve *T* as in Eq. 5. Within group, the tuning amplitude for each neuron is varied. To this end, we sample a vector of 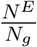 random number from a normal distribution with *μ* = 0 and *σ* = 0.05. We then shuffle this vector for each group and add a single value to the synaptic tuning curve of each neuron in the group.

The excitatory and inhibitory CA1 neurons also receive uniform external currents, 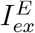 and 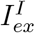 respectively.

#### Excitatory plasticity

The excitatory synaptic weights between CA3 and CA1 excitatory neurons follow classical Hebbian plasticity implemented using a symmetrical spike-timing dependent learning rule. A synaptic trace *x_i_* assigned to each neuron *i* and follows:

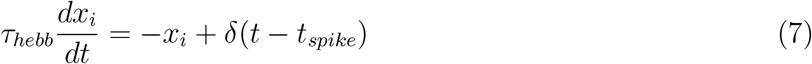

where *τ_hebb_* is the time constant of the learning window and *t_spike_* is the spike time. We also include a homeostatic term which takes into account the sum of all synaptic weights onto the postsynaptic neuron. The synaptic weight *w_ji_* from the presynaptic neuron *j* to the postsynaptic neuron *i* is updated following:

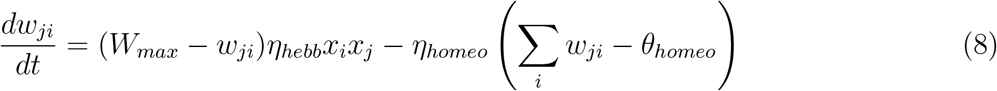

where *W_max_* is the maximum excitatory synaptic weight, *η_hebb_* is the learning rate of the Hebbian term, *η_homeo_* is the learning rate of the homeostatic term and *θ_homeo_* is the homeostatic target.

### Recurrent Network Model

We simulate a recurrent network consisting of *N_E_* excitatory and *N_I_* inhibitory neurons. The excitatory neurons are randomly connected with a probability *p_EE_* = 0.1, in line with experiments (Harris & Shepherd 2015). The inhibitory neurons are randomly connected to the excitatory neurons with a probability *p_IE_* = *p_EI_* = 0.2. Due to the small size of the network, the *in* degree for each excitatory and inhibitory neuron is kept the same. Thus each excitatory neuron has *N_E_* × *p_EE_* excitatory inputs and 4 *N_I_* × *p_IE_* inhibitory recurrent synaptic inputs. Furthermore, each inhibitory neuron receives *N_E_* × *p_EI_* excitatory inputs. Due to the small network size, inhibitory to inhibitory neuron connections are omitted (*p_II_* = 0). The values for all other synaptic weights are sampled from a log normal distribution, in line with experiments (Song et al. 2005, Loewenstein et al. 2011), with *μ* = 1.0 and *σ* = 0.1.

The initial values of the inhibitory synaptic weights are set to ten times weaker than the excitatory weights. Furthermore, the inhibitory synaptic weight update Δ*w_ji_* has a multiplicative weight dependence, such that:

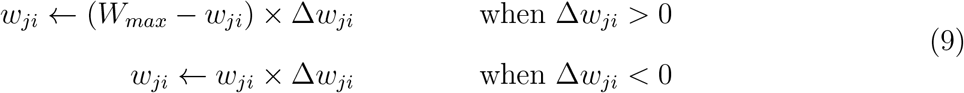

where *W_max_* is the is the maximum inhibitory synaptic weight. The inhibitory learning is turned off during the first 15 seconds of the simulation.

#### Random neuron input generation

Each neuron in the network is randomly connected with a probability of *p_ex_* = 0.2 to a subset of the *N_ex_* external inputs, which are modelled as Poisson process with a mean firing rate *F*. To generate sufficient network activity, the synaptic weights from the external inputs to the excitatory and inhibitory neurons are 2.5 times stronger than the mean recurrent excitatory weight. For the simulation in Fig 2.A, the values of the synaptic weights from the external inputs to two excitatory neurons are increased by 50%.

#### Memory protocol

The same network as above is used in the simulation. After the network activity stabilises (600 s), the recurrent synapses between a selected subset of *N^E^* (12/80) are increased by a factor of 5. After the network reaches steady state again (1200 s), the values of the synaptic weights from the 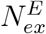 to two neurons in the assembly are increased by 50%.

#### Inhibitory spike-timing dependent plasticity (iSTDP)

With iSTDP (Vogels et al. 2011), the synaptic weight *W_ji_* from presynaptic inhibitory neuron *i* and the postsynaptic excitatory neuron *j* follows:

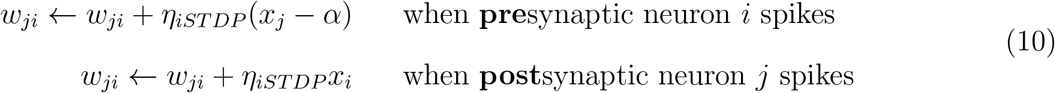

where *η_STDP_* is the learning rate, *α* is the depression factor and *x_i/j_* is the neuron synaptic trace, defined as:

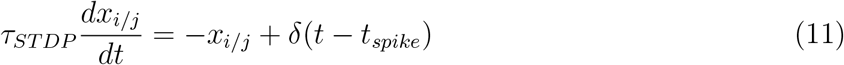

where *τ_iSTDP_* is the learning window time constant and *t_spike_* is the spike time.

## Code

In all the simulations, we use Euler integration with step size Δ*t* of 1 ms. The Python code will be made freely available after publication (GitHub and ModelDB) and is given to the reviewer.

## Acknowledgements

This work was supported by BBSRC BB/N013956/1, BB/N019008/1, Wellcome Trust 200790/Z/16/Z, Simons Foundation 564408 and EPSRC EP/R035806/1 and a Wellcome Trust PhD award to KK.

## Parameters summary

Parameters common across both simulated networks.

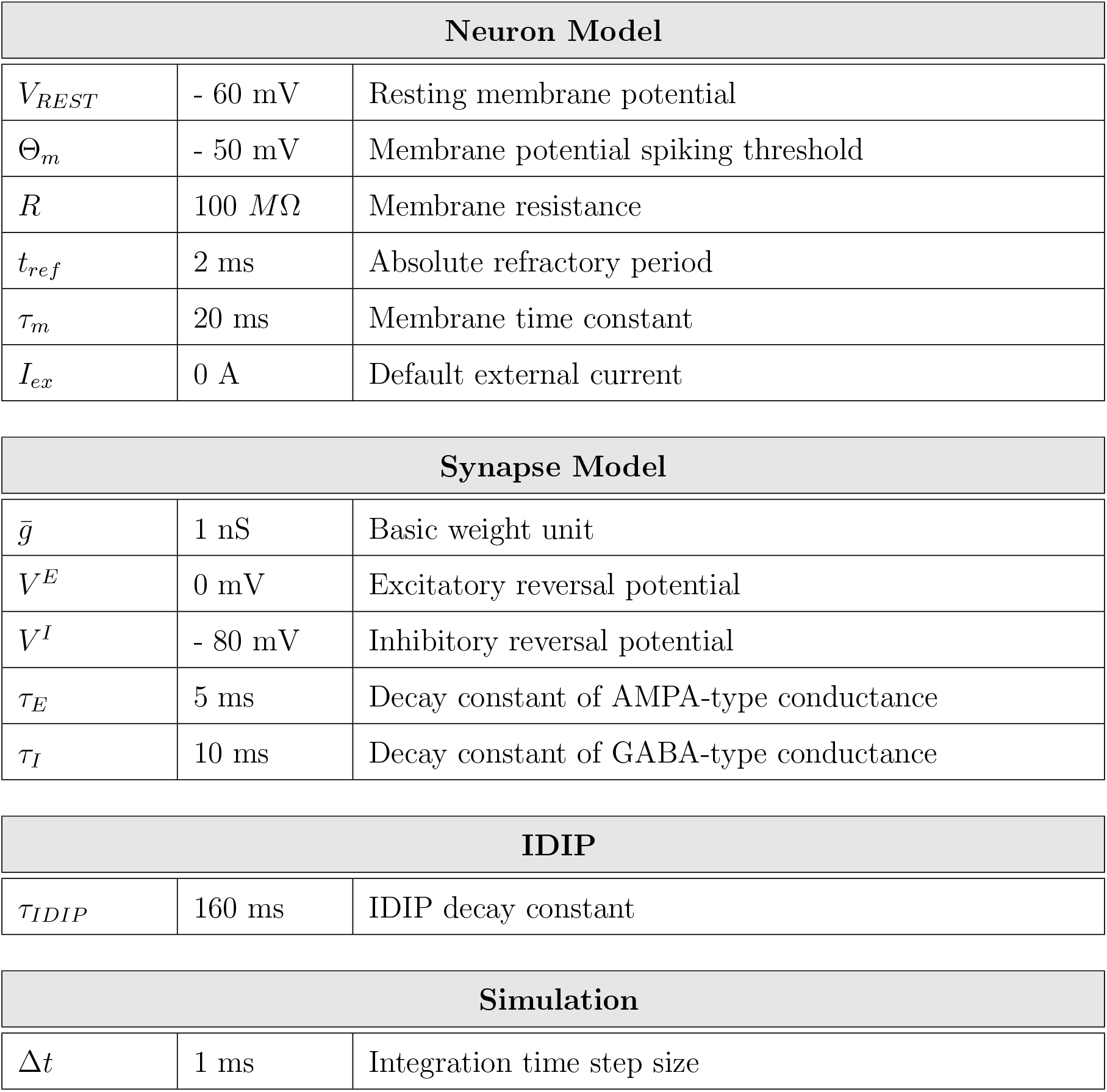

| Neuron Model |  |  |
| --- | --- | --- |
| $V_{REST}$ | - 60 mV | Resting membrane potential |
| $\Theta_m$ | - 50 mV | Membrane potential spiking threshold |
| $R$ | 100 $M\Omega$ | Membrane resistance |
| $t_{ref}$ | 2 ms | Absolute refractory period |
| $\tau_m$ | 20 ms | Membrane time constant |
| $I_{ex}$ | 0 A | Default external current |

| Synapse Model |  |  |
| --- | --- | --- |
| $\bar{g}$ | 1 nS | Basic weight unit |
| $V^E$ | 0 mV | Excitatory reversal potential |
| $V^I$ | - 80 mV | Inhibitory reversal potential |
| $\tau_E$ | 5 ms | Decay constant of AMPA-type conductance |
| $\tau_I$ | 10 ms | Decay constant of GABA-type conductance |

| IDIP |  |  |
| --- | --- | --- |
| $\tau_{IDIP}$ | 160 ms | IDIP decay constant |

| Simulation |  |  |
| --- | --- | --- |
| $\Delta t$ | 1 ms | Integration time step size |

| Hippocampal Network Model |  |  |  |
| --- | --- | --- | --- |
| **CA3 neurons** | $N_{CA3}$ | 10 | Size of the CA3 excitatory population |
| **CA1 neurons** | $N_E$ | 100 | Size of the CA1 excitatory population |
| | $N_I$ | 20 | Size of the CA1 inhibitory population |
| | $N_g$ | 10 | Number of excitatory neuron groups |
| **Position modulated input** | $N_{pc}$ | 10 | Number of place fields |
| | $\sigma_{pc}$ | 6.54 a.u. | Tuning width |
| | $A_c$ | 200 pA | Current amplitude |
| **External current** | $I_{ex}^E$ | 0.01 pA | Current to the excitatory neurons |
| | $I_{ex}^I$ | 0.5 pA | Current to the inhibitory neurons |
| **Synaptic weights** | $W_{EI}$ | 2.0 | E to I synaptic weight |
| | $W_{IE}$ | 0.1 | Initial I to E synaptic weight |
| **IDIP** | $\eta_{IDIP}$ | 10 | IDIP learning rate |
| | $\theta_{in}$ | 200 nS | Inhibitory neuron target input |
| **Excitatory plasticity** | $\tau_{hebb}$ | 20 ms | Learning window time constant |
| | $W_{max}$ | 1.5 | Maximum E to E synaptic weight |
| | $\eta_{hebb}$ | $0.01 \text{ s}^{-1}$ | Excitatory Hebbian term learning rate |
| | $\eta_{homeo}$ | $0.001 \text{ s}^{-1}$ | Excitatory homeostatic term learning rate |
| | $\theta_{homeo}$ | 5.2 | Homeostatic target sum of weights |

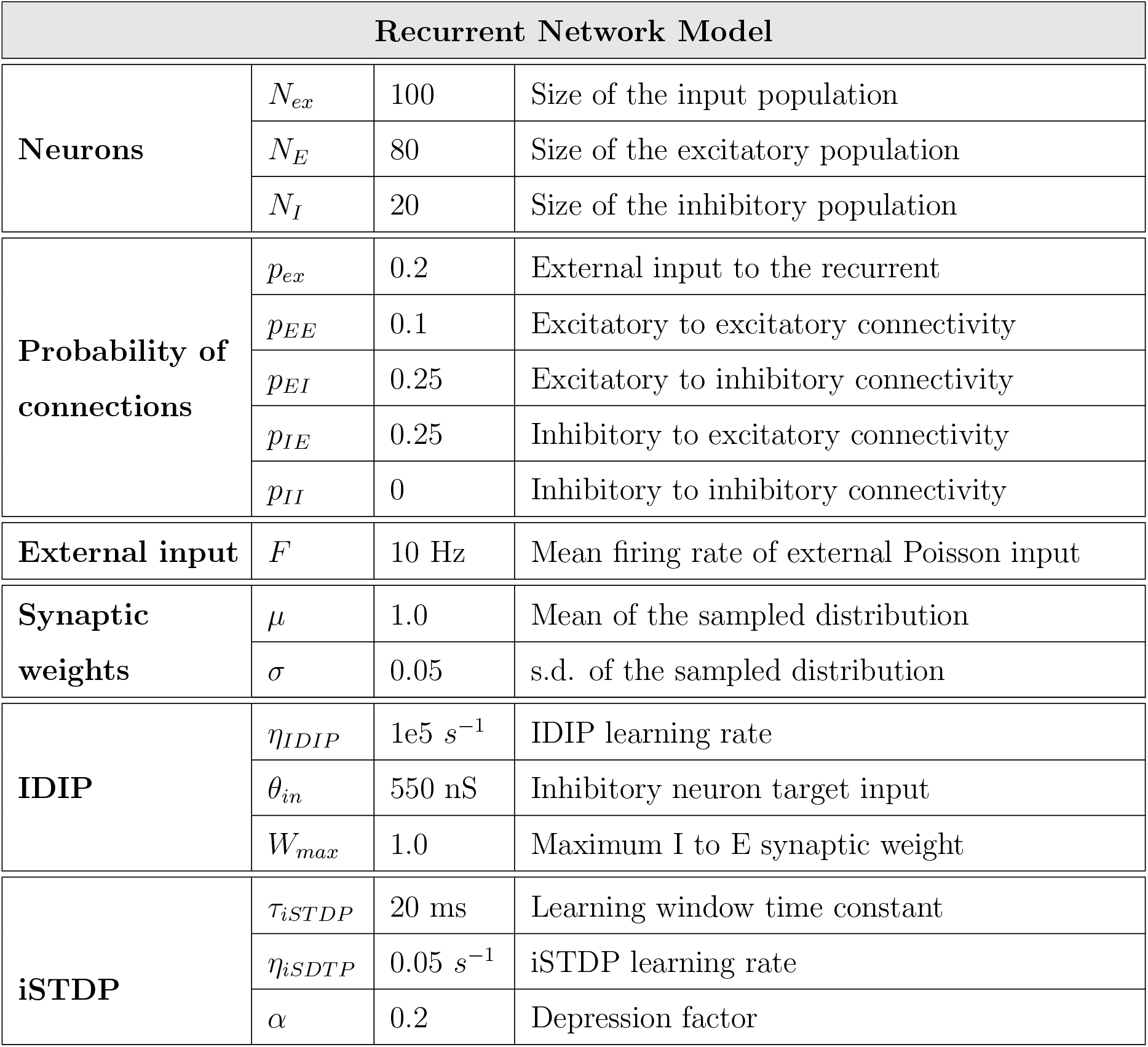

## Notes

### Competing Interest Statement

The authors have declared no competing interest.

