## Supplementary Figures for "Network-centered homeostasis through inhibition maintains hippocampal spatial map and cortical circuit function"

A

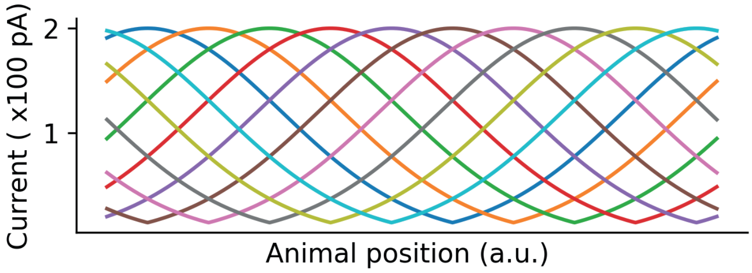


B

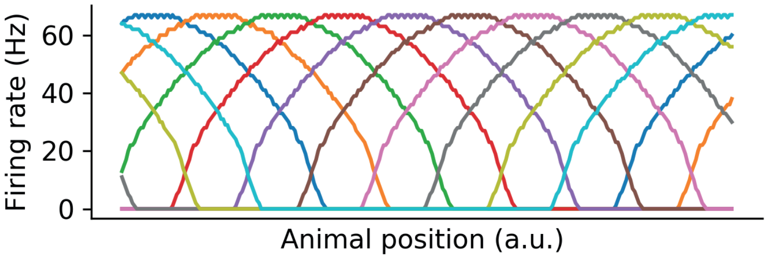


**Supplementary Figure 1.** **A** The external current supplied to the CA3 neurons. Different colours indicate different place fields. **B** The CA3 neuron firing rates. Different colours indicate different CA3 neurons tuned to different place fields, as in **A**.

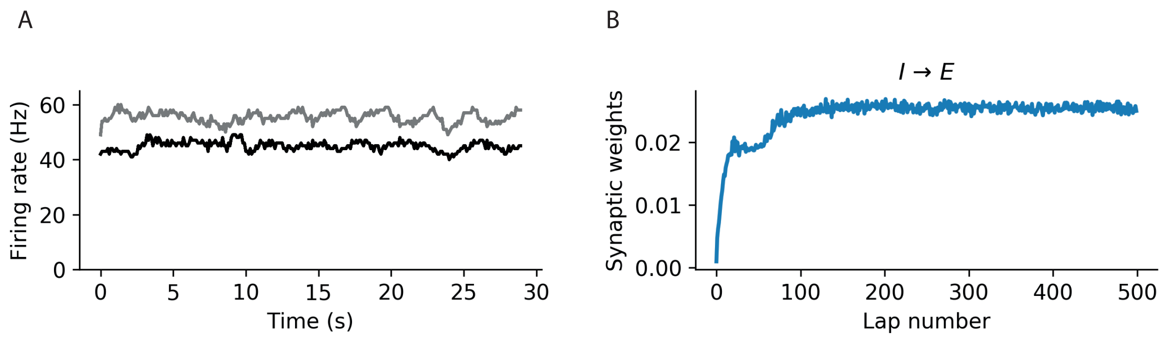


**Supplementary Figure 2.** **A** Firing rate of the CA1 inhibitory neurons in the first (grey) and last (black) lap of the simulation. **B** CA1 inhibitory synaptic weights as a function of time.

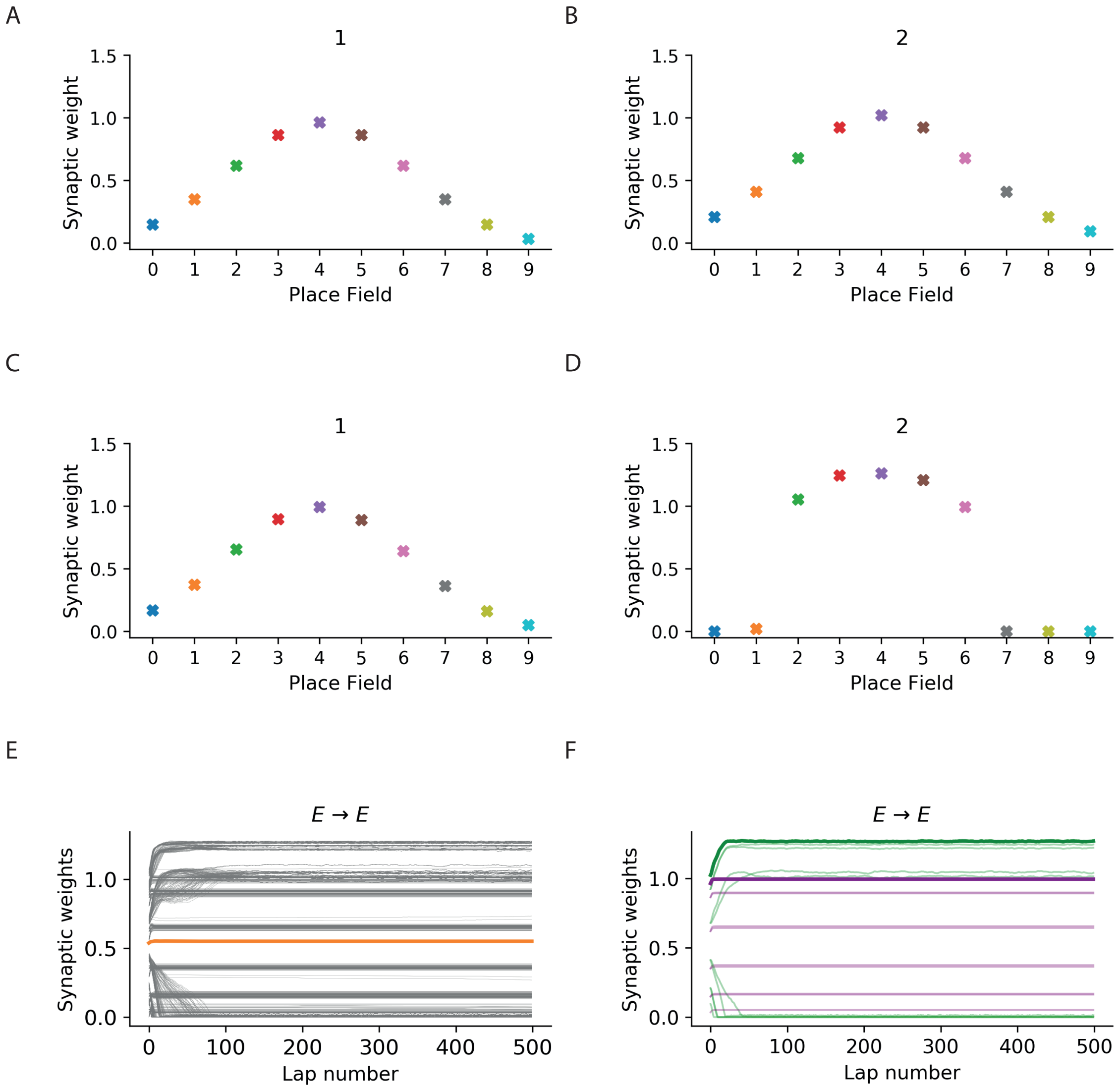


**Supplementary Figure 3.** **A** The initial CA3 to CA1 synaptic weights of neuron 1, as in Fig 1.C. Different colours indicate synapses from CA3 neurons with different place fields, as in S1. **B** The initial CA3 to CA1 synaptic weights of neuron 2, as in Figure 1.C. **C** The final CA3 to CA1 synaptic weights of neuron 1, as in Figure 1.F. **D** The final CA3 to CA1 synaptic weights of neuron 2, as in Figure 1.F. **E** CA3 to CA1 synaptic weights as a function of time. Grey lines indicate individual weights and the orange line denotes the mean. **F** CA3 to CA1 synaptic weights of an example place (green) and silent (purple) cell as a function of time. Pale lines indicate individual weights and the bold lines denote the mean.

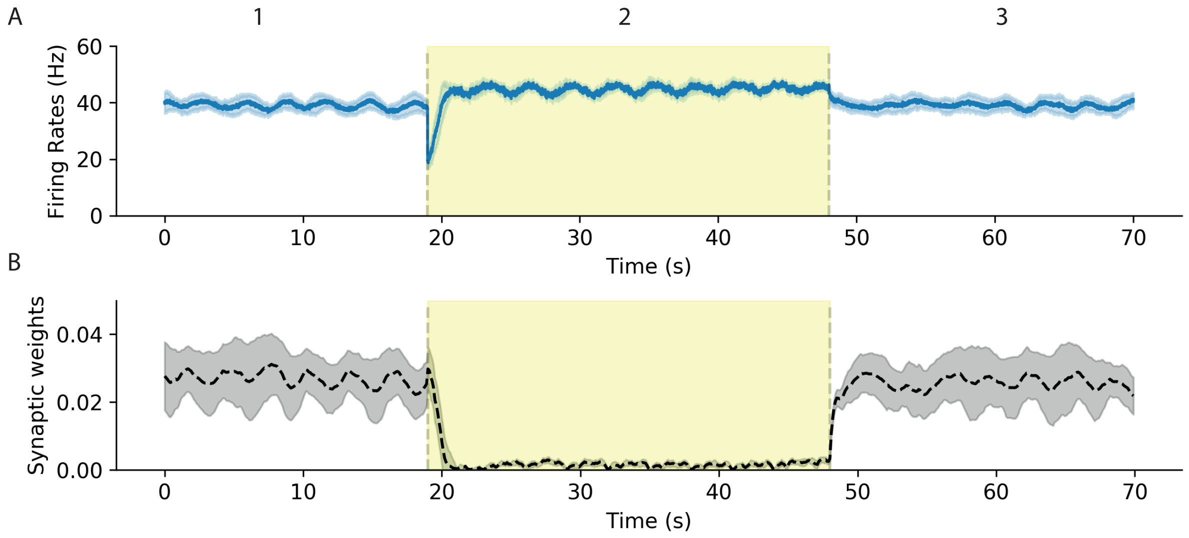


**Supplementary Figure 4.** **A** Evolution of the inhibitory firing rates during the silencing protocol as described in Fig 2.A. **B** Evolution of the inhibitory synaptic weights during the silencing protocol as described in Fig 2.A.

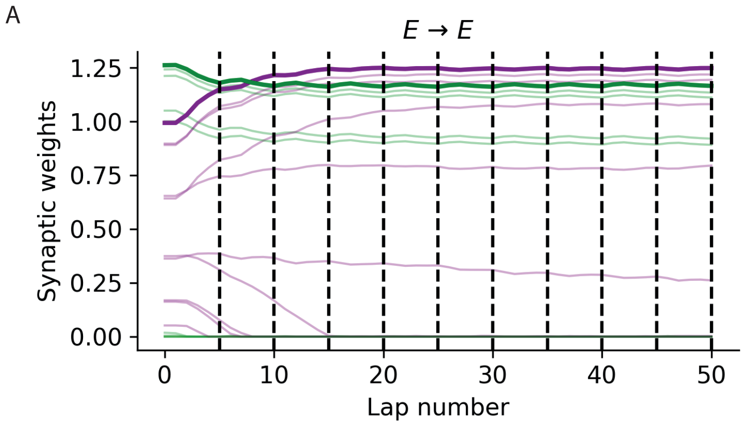


**Supplementary Figure 5. A** Evolution of the synaptic weights from CA3 to CA1 for an example place (green) and silent (purple) cell during the consolidation protocol as described in Fig 2.I (mean in bold). Dashed vertical lines mark testing laps.

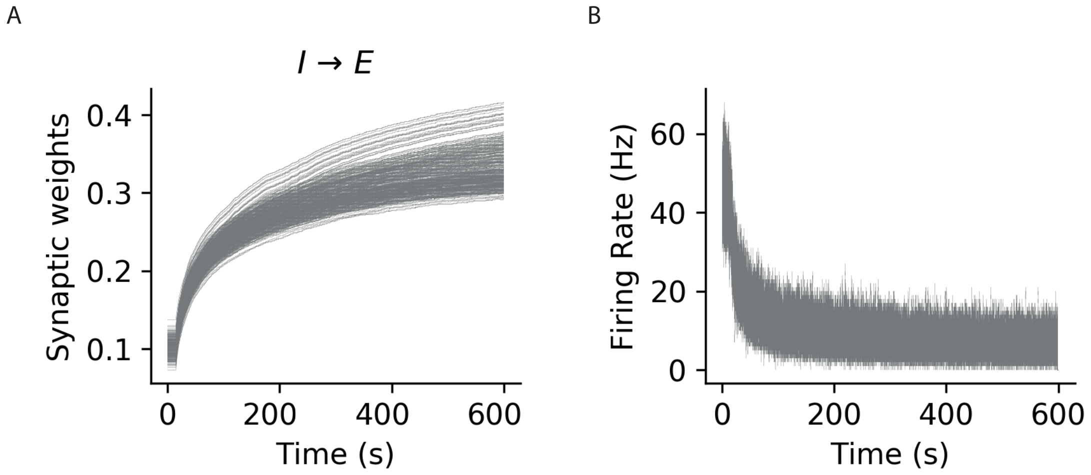


**Supplementary Figure 6.** Recurrent network. **A** All the inhibitory synaptic weights. **B** All the excitatory firing rates.

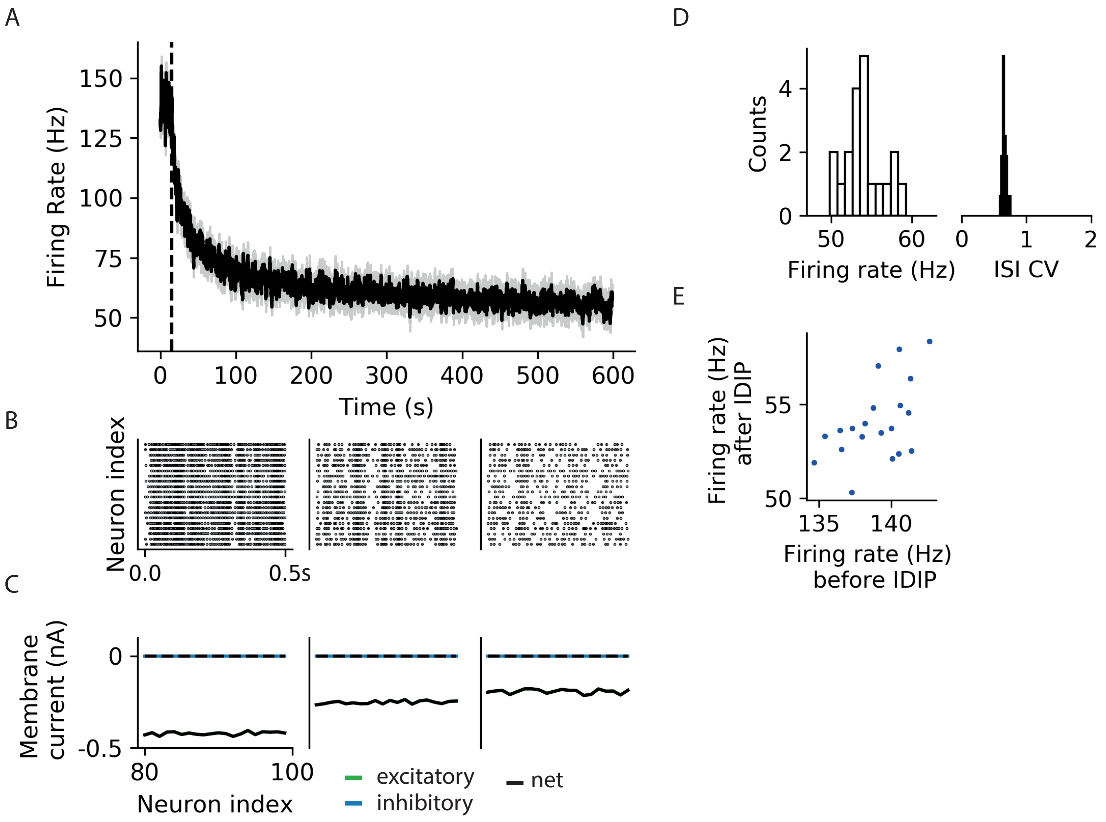


**Supplementary Figure 7.** **A-D** Same as Fig 3.B-C,E-F but for the inhibitory neurons. **E** The initial (x-axis) versus final (y-axis) inhibitory firing rates.

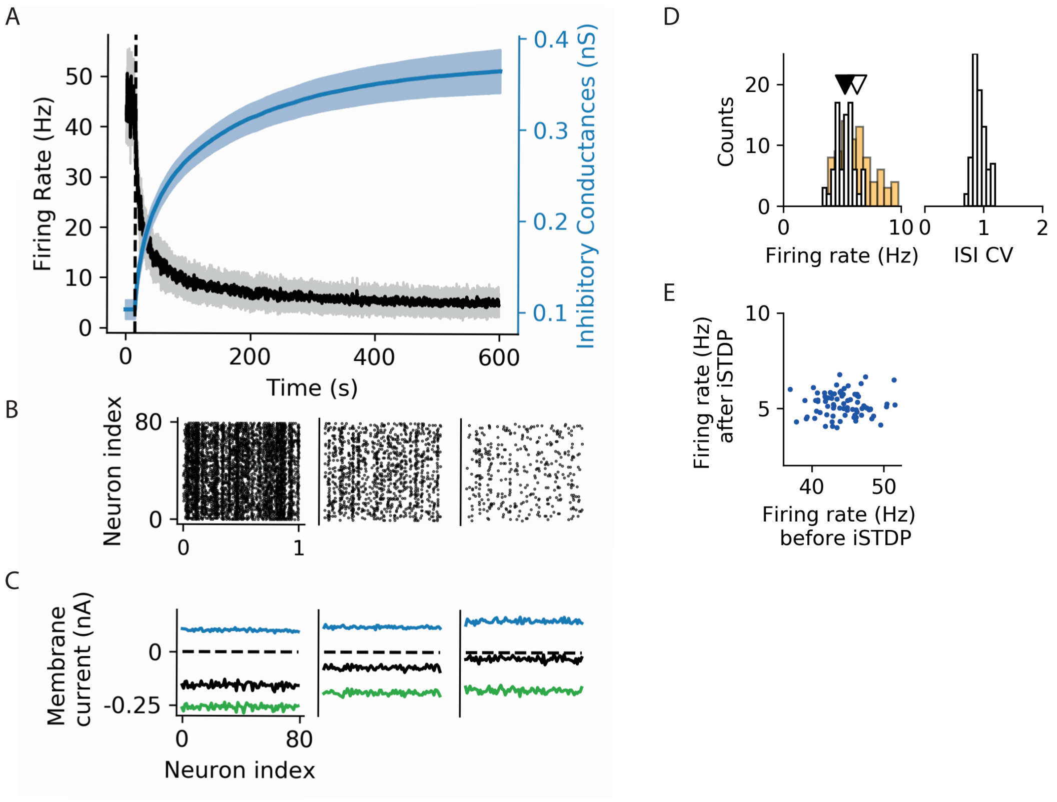


**Supplementary Figure 8.** **A-D** Same as Fig 3.B-C,E-F but with networks following iSTDP (**?**). **D** The pale orange indicates the distribution of firing rates with IDIP in the same network. The mean network firing rates are indicated with triangles for iSTSP (black) and IDIP (white). **E** The initial (x-axis) versus final (y-axis) inhibitory firing rates.

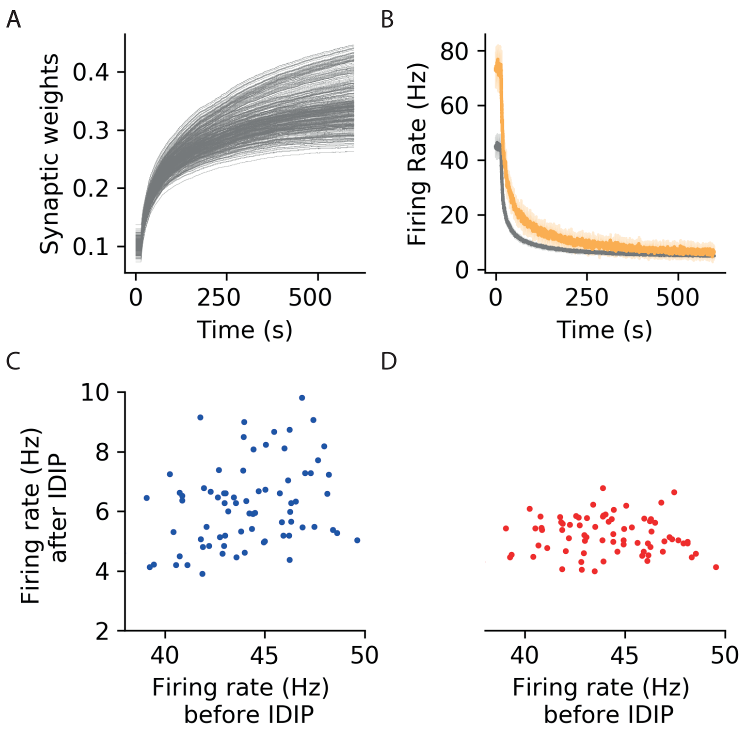


**Supplementary Figure 9.** **A** The evolution of all the inhibitory synaptic weights while some neurons receive increased input, as in Fig 4.A . **B** Same as Fig 4.B but in a network following iSTDP (**?**). **C** The excitatory firing rate before (x-axis) versus after (y-axis) inhibitory learning in the network following IDIP. **D** The excitatory firing rate before (x-axis) versus after (y-axis) inhibitory learning in the network following iSTDP (**?**).

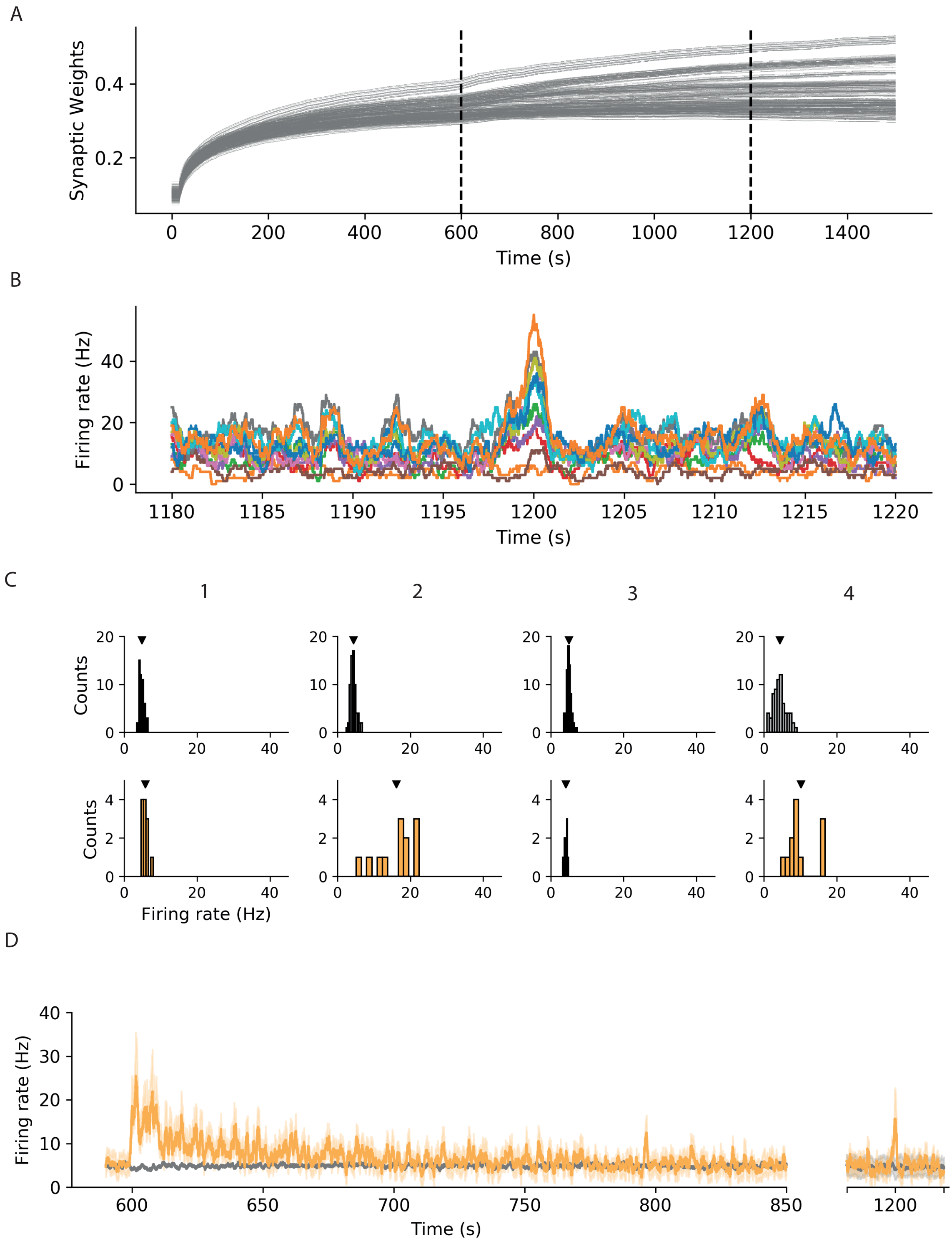


**Supplementary Figure 10. A** All the inhibitory synaptic weights during the associative memory task (Fig 4.F). The dashed vertical lines indicate memory encoding and recall respectively. **B** Close-up of all the firing rates in the assembly during recall. **C-D** Same as Fig 4.G-H but in a network following iSTDP (**?**).

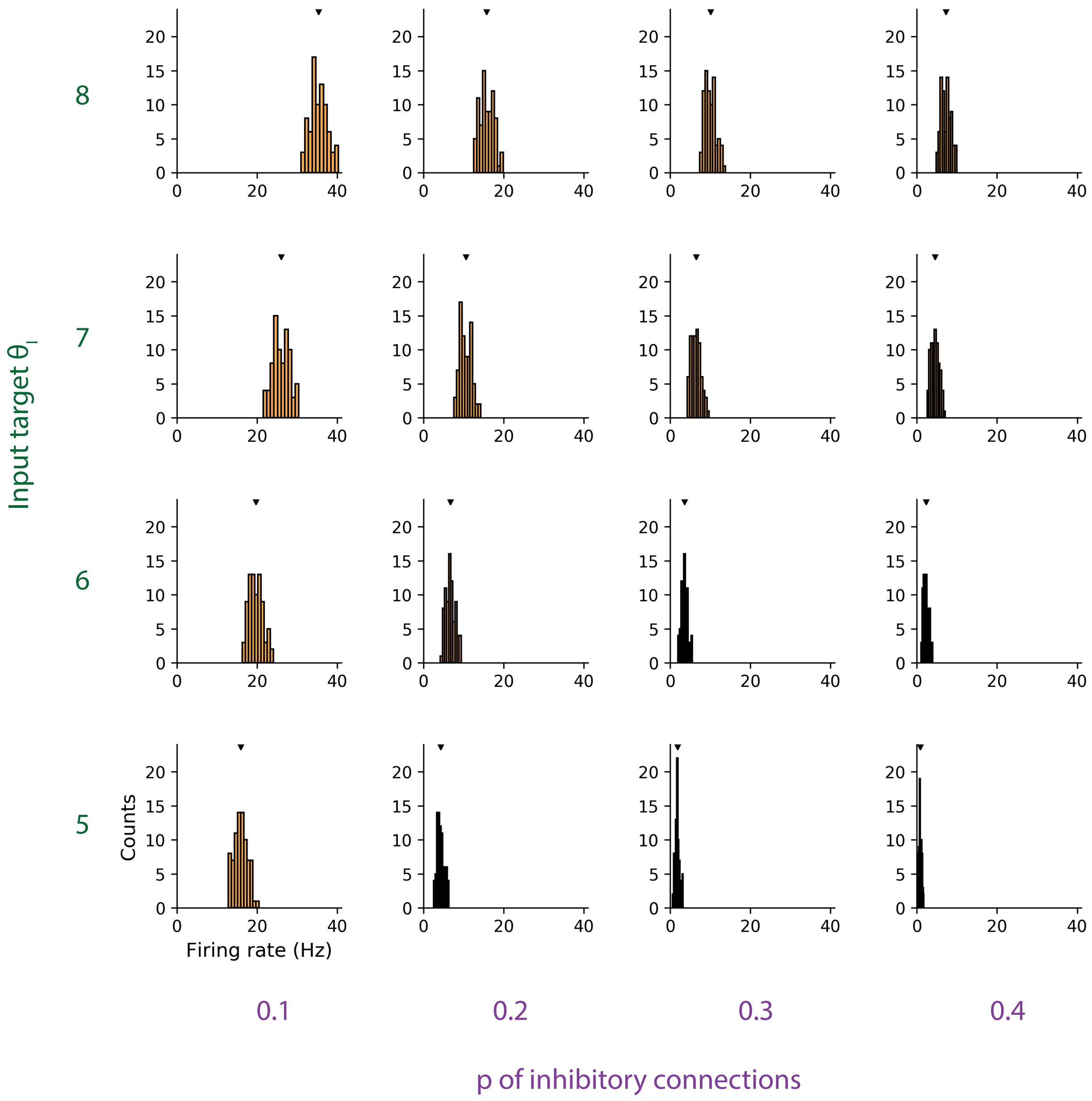


**Supplementary Figure 11.** Same as Figure 3.D but distribution of firing rates as a function of different values of inhibitory probability of connections (horizontal axis) and input target $\theta_i$ (vertical axis). The means are denoted with black triangles.

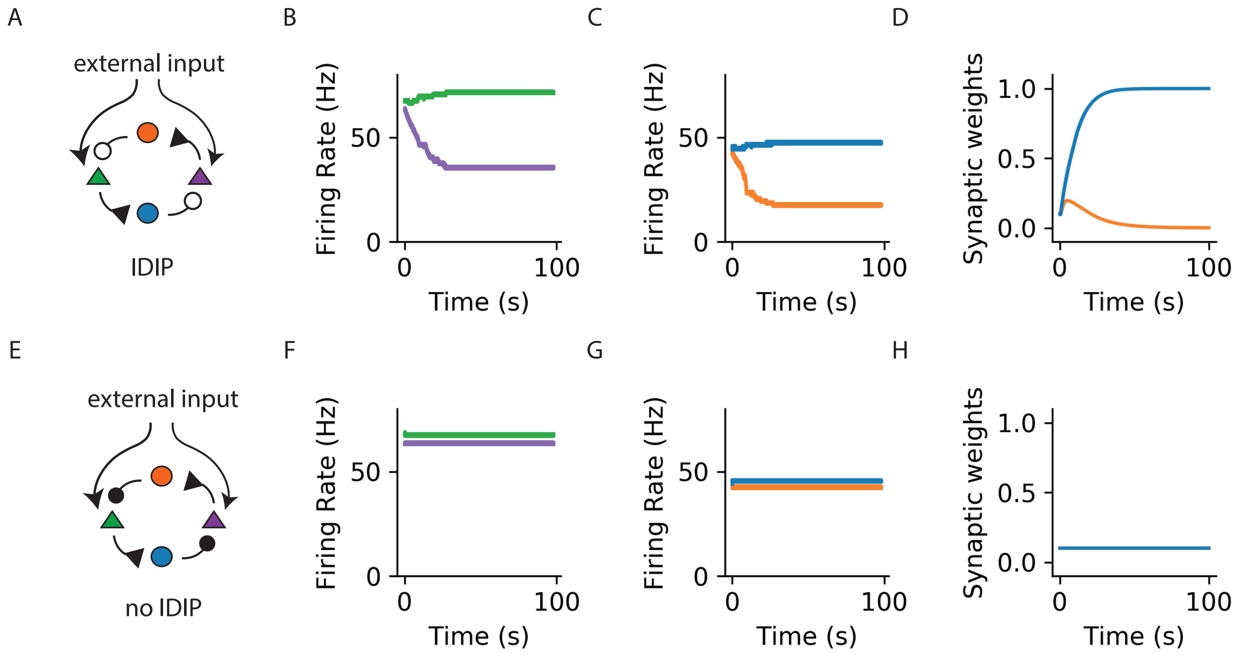


**Supplementary Figure 12. A-D** In the limit of very sparse networks, IDIP may not act as a homeostatic mechanism. **A** Network diagram. Two excitatory (triangles) and two inhibitory (circles) neurons are simulated. One of the excitatory neurons (green) receives a slightly (5%) stronger external input. Thus one inhibitory neuron (blue) receives synaptic input from the excitatory neuron that is more active and projects onto the other excitatory neuron (purple). The other inhibitory neuron (orange) receives synaptic input from the less active excitatory neuron and projects onto the more active excitatory neuorn. **B** Evolution of neuronal firing rates for the excitatory cells, color coded as in **A**. The initially small activity difference is amplified by inhibitory plasticity. **C** Evolution of neuronal firing rates for the inhibitory cells, color coded as in **A**. **D** Evolution of the inhibitory (I to E) synaptic weights. **E-H** The same as in **A-D** but without any plasticity.
